# V-pipe: a computational pipeline for assessing viral genetic diversity from high-throughput sequencing data

**DOI:** 10.1101/2020.06.09.142919

**Authors:** Susana Posada-Céspedes, David Seifert, Ivan Topolsky, Karin J. Metzner, Niko Beerenwinkel

## Abstract

High-throughput sequencing technologies are used increasingly, not only in viral genomics research but also in clinical surveillance and diagnostics. These technologies facilitate the assessment of the genetic diversity in intra-host virus populations, which affects transmission, virulence, and pathogenesis of viral infections. However, there are two major challenges in analysing viral diversity. First, amplification and sequencing errors confound the identification of true biological variants, and second, the large data volumes represent computational limitations. To support viral high-throughput sequencing studies, we developed V-pipe, a bioinformatics pipeline combining various state-of-the-art statistical models and computational tools for automated end-to-end analyses of raw sequencing reads. V-pipe supports quality control, read mapping and alignment, low-frequency mutation calling, and inference of viral haplotypes. For generating high-quality read alignments, we developed a novel method, called *ngshmmalign*, based on profile hidden Markov models and tailored to small and highly diverse viral genomes. V-pipe also includes benchmarking functionality providing a standardized environment for comparative evaluations of different pipeline configurations. We demonstrate this capability by assessing the impact of three different read aligners (Bowtie 2, BWA MEM, ngshmmalign) and two different variant callers (LoFreq, ShoRAH) on the performance of calling single-nucleotide variants in intra-host virus populations. V-pipe supports various pipeline configurations and is implemented in a modular fashion to facilitate adaptations to the continuously changing technology landscape. V-pipe is freely available at https://github.com/cbg-ethz/V-pipe.

## 1 Introduction

RNA viruses are regarded as important models in evolutionary biology. They exhibit short generation times, and their mutation rates are higher than those of any other living organism (Duffy *et al*., 2008). As a result, they evolve rapidly and often exist within their host as a collection of distinct, yet genetically related, viral strains, which are constantly adapting to respond to challenging environments (Lauring and Andino, 2010). The existence of a heterogeneous mixture of viral strains, often referred to as a viral quasispecies (Domingo *et al*., 2005), impacts viral transmission, virulence, and pathogenesis (Vignuzzi *et al*., 2006; Tsibris *et al*., 2009; Rozera *et al*., 2014). Moreover, the emergence and accumulation of mutations during the course of a viral infection limits the success of antiviral therapies, for example, in Human Immunodeficiency Virus (HIV) (Wensing *et al*., 2019), Hepatitis C Virus (HCV) (Hiraga *et al*., 2011; Wyles and Luetkemeyer, 2017), or influenza infections (Hu *et al*., 2013). Therefore, analysing the genetic structure of virus populations is highly relevant in the context of immune escape (Nowak *et al*., 1991; Kuroda *et al*., 2010), as well as resistance to vaccines (Gaschen *et al*., 2002) and antiviral drugs (Mason *et al*., 2018).

High-throughput sequencing (HTS) technologies offer the possibility of sequencing many of the viral strains, as opposed to determining only the consensus sequence of the population by Sanger sequencing, which neglects minor variants (Zagordi *et al*., 2010b). Indeed, variants with a relative frequency below 20% cannot be reliably identified with Sanger sequencing (Schuurman *et al*., 1999; Wang *et al*., 2007). Although low-frequency variants may be clinically relevant, e.g., in the context of drug resistance or vaccine response (Johnson *et al*., 2008; Henn *et al*., 2012; Vandenhende *et al*., 2014; Cozzi-Lepri *et al*., 2015; Kyeyune *et al*., 2016; Mbunkah *et al*., 2020), Sanger sequencing remains the gold standard in many clinical applications.

HTS technologies have opened up new possibilities for in-depth characterization of the genetic diversity of virus samples (Barzon *et al*., 2013; Capobianchi *et al*., 2013). In order to incorporate the use of these technologies into research and clinical diagnostics, standardized procedures for data generation and analysis are required. However, analysing viral HTS data is complicated by large volumes of data, short length of the sequencing reads, and the high amplification and sequencing error rates relative to the expected intra-host viral diversity (Goodwin *et al*., 2016). Therefore, statistical and computational challenges remain in disentangling true biological variation from technical errors, especially for low-frequency variants (Beerenwinkel *et al*., 2012).

Several methods have been proposed for studying genetic diversity in virus populations. Genetic variation has been addressed at the level of (*i*) single nucleotide variants (SNVs) (Wang *et al*., 2007; Wilm *et al*., 2012; Macalalad *et al*., 2012; Flaherty *et al*., 2012; Yang *et al*., 2013; McElroy *et al*., 2013), (*ii*) short variant sequences, or local haplotypes, constrained by the read length (Zagordi *et al*., 2010a), and (*iii*) long-range viral haplotypes by phasing mutations over distances larger than the read length (Prosperi and Salemi, 2012; Töpfer *et al*., 2013, 2014; Prabhakaran *et al*., 2014; Baaijens *et al*., 2017; Artyomenko *et al*., 2017; Leviyang *et al*., 2017). Some of these tools appear no longer actively maintained or limited in terms of robustness and usability. Also, the analysis of HTS data involve additional steps implemented by separate tools, e.g., quality control and read alignment. A prerequisite for incorporating such HTS data into routine diagnostics is to standardize the processing steps end-to-end from raw data input to final output.

In recent years, several bioinformatics pipelines have been developed for different viral HTS data analysis tasks. Many workflows have been designed for virus discovery and metagenomics applications (Ho and Tzanetakis, 2014; Naccache *et al*., 2014; Li *et al*., 2016; Zhao *et al*., 2017; Zheng *et al*., 2017; Maarala *et al*., 2017). Other pipelines focus on either (*i*) the construction of the consensus sequence via reference-guided assembly, such as VirAmp (Wan *et al*., 2015) and shiver (Wymant *et al*., 2018), (*ii*) the identification of SNVs, such as hivmmer (Howison *et al*., 2018), HyDRA (Taylor *et al*., 2019) and MinVar (Huber *et al*., 2017), or (*iii*) the reconstruction of viral haplotypes, such as VGA (Mangul *et al*., 2014) and ViQuaS (Jayasundara *et al*., 2015). Often these tools either target specific viruses (Huber *et al*., 2017; Wymant *et al*., 2018; Howison *et al*., 2018; Taylor *et al*., 2019) or correspond to proof-of-concept implementations with limited support for broader applications. For HIV drug resistance testing, Lee *et al*. (2020) have recently evaluated various bioinformatics pipelines for this specific application.

An important step in inferring viral genetic diversity is the alignment of HTS reads. There are two general strategies for read alignment, namely reference-based approaches and *de novo* assembly. A limitation of the former is the introduction of biases due to differences in sequence similarity between the reference genome and viral haplotypes (Archer *et al*., 2010). In addition to potential reference biases, placement of gaps remains a challenge in aligning reads to a reference sequence. Reads containing insertions or deletions (indels) are less likely to be accurately aligned (Posada-Céspedes *et al*., 2017). On the other hand, limitations of *de novo* assembly include increased sensitivity to chimeric reads leading to erroneous contigs, and a segmented coverage of the genome by the assembled contigs (Wymant *et al*., 2018).

To address these limitations, we have developed a novel read alignment tool, called *ngshmmalign*. The aligner borrows ideas from the alignment of protein families to align HTS reads from small and highly diverse genomes. Ngshmmalign models the multiple read alignment as a profile hidden Markov model (HMM) and aligns all reads to this profile. By doing so, ngshmmalign accounts for local heterogeneity, including structural variations.

To support reproducible viral genomics studies for basic research and clinical diagnostics, we have developed V-pipe, a flexible bioinformatics pipeline integrating several tools for analysing viral HTS data. V-pipe allows for assessing viral diversity on all three genomic scales described above: SNVs, short variant sequences, and long-range haplotypes. To provide the flexibility required for adapting to future methodological and technological developments, V-pipe provides a modular and extensible framework which facilitates the introduction of new tools. V-pipe also supports the comparative assessment of different workflows in a standardized environment. To this end, it contains modules to generate synthetic data and to assess the accuracy of the computational inference. We demonstrate the benchmark capabilities of V-pipe by assessing the impact of different read aligners and variant callers on the accuracy of SNV calling. We validate the pipeline using sequencing data from a control sample composed of five well-defined HIV-1 strains (Di Giallonardo *et al*., 2014), and to demonstrate its applicability, we process 92 HIV-1 whole-genome sequencing samples from 11 patients previously reported in Zanini *et al*. (2015).

## 2 Methods

We first summarise the core components of V-pipe. We then present the novel read alignment tool ngshmmalign and explain the benchmarking functionalities of the pipeline. Lastly, we describe the simulation setup and the control sequencing data sets used to assess the performance of different components of V-pipe.

### 2.1 Computational pipeline

V-pipe uses the Snakemake workflow management system (Köster and Rahmann, 2012), which enables the well-controlled and scalable execution of the pipeline in local as well as in high-performance computing environments. The pipeline integrates various open-source software packages developed for analysing virus samples. For deployment, we provide Conda (https://conda.io) environments to automatically download and install the required tools. The environments include all dependencies and versions which is key for full reproducibility of the analysis settings. As input, V-pipe requires the raw sequencing data in FASTQ format, a reference sequence in FASTA format, and a configuration file containing user-defined options. V-pipe supports both single-end and paired-end FASTQ files. In short, the analysis workflow implemented by V-pipe involves the following main steps: (*i*) quality control, (*ii*) reference-guided mapping and alignment of sequencing reads, and (*iii*) identification of SNVs and reconstruction of viral haplotypes (Fig. 1).

**Figure 1.**
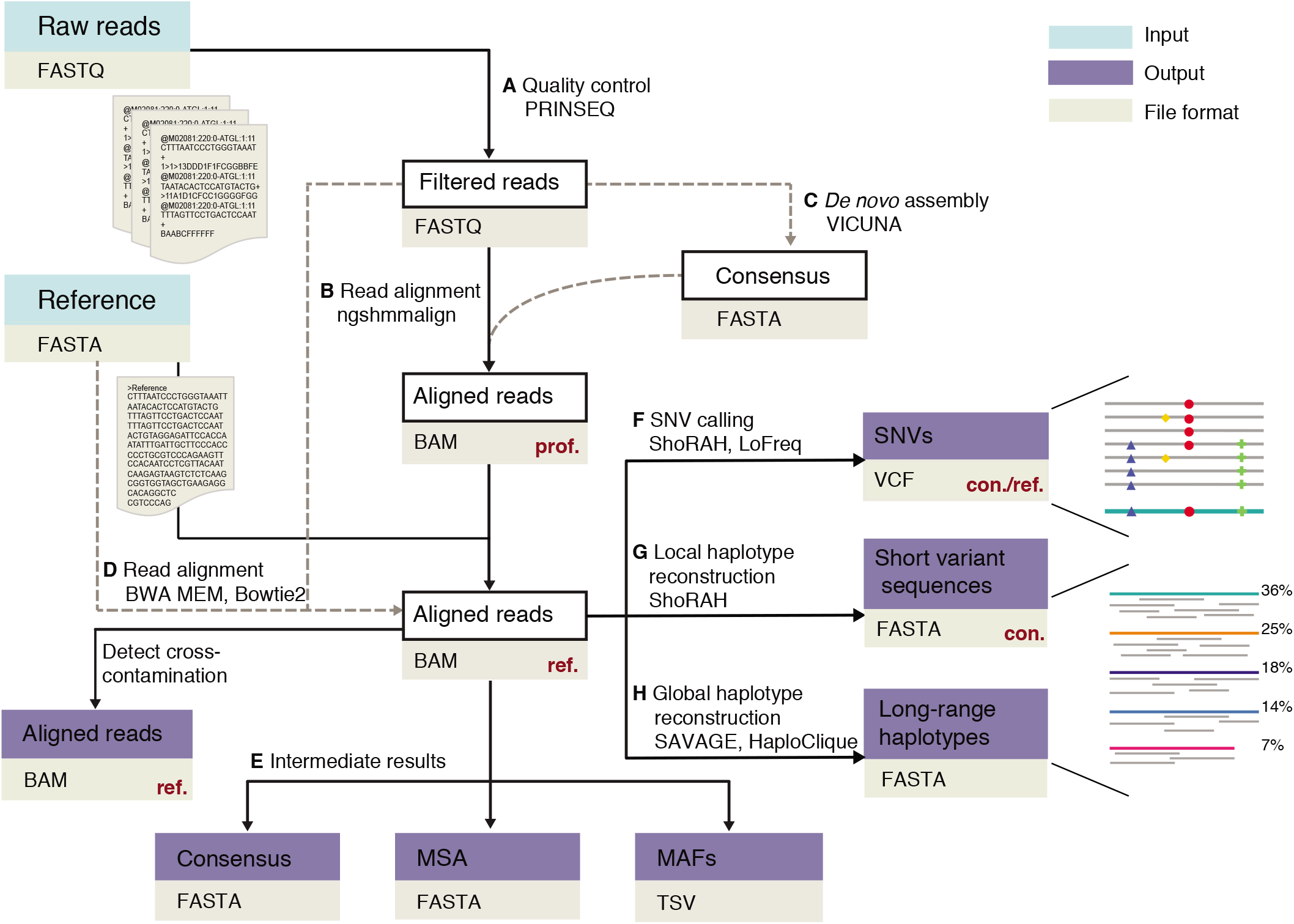
Workflow of V-pipe for the analysis of viral HTS data. As input, the pipeline requires raw sequencing data (FASTQ format) and a reference sequence (FASTA format), which defines the indexing frame for the reporting of variants. **A** For quality control, low-quality bases are removed from both termini. **B** Reads are aligned employing a reference-guided approach using ngshmmalign. **C** For the read alignment, the reference sequence may be provided or it can be built *de novo* from the read data. **D** Alternatively, reads can be aligned using BWA MEM or Bowtie2. **E** Intermediate results are provided in the form of a consensus sequence per sample, a multiple sequence alignment of all consensus sequences, and the minor allele frequencies (MAFs) for all samples and all loci. **F** Single-nucleotide variants (SNVs) are identified, and **G** local and **H** global haplotypes are reconstructed. Whenever applicable, the pertinent reporting frame (con.: consensus, prof.: profile, ref: reference) is indicated in red. Dotted lines indicate alternative processing steps included in V-pipe.

Relatively high error rates are a limitation of HTS technologies. Error sources include the reverse transcription of viral RNA into cDNA, the amplification of the material by multiple cycles of PCR, and the sequencing process itself. To avoid propagating biases to the downstream analysis steps, it is imperative to include quality checks and filtering steps. To this end, we include FastQC (Andrews, 2019) to provide quality control reports and PRINSEQ (Schmieder and Edwards, 2011) for removing low-quality or ambiguous bases from both termini of reads.

For aligning sequencing reads, we developed a novel read aligner called ngshmmalign which is described below. This aligner requires as input a reference sequence to approximately locate reads. The reference sequence can be provided by the user or constructed *de novo* using the VICUNA software. We also include two alternative aligners, namely BWA MEM (Li, 2013) and Bowtie2 (Langmead and Salzberg, 2012), to choose from according to particular needs and computational resources. In addition to the alignment file, ngshmmalign outputs two types of consensus sequences constructed from the aligned reads, namely by (*i*) using a majority vote at each position, and (*ii*) incorporating ambiguous bases. V-pipe produces these consensus sequences for the alternative aligners. We use lowercase characters in the consensus sequences to mark positions with read counts below 50 reads, by default. To report ambiguous bases, every base with a relative frequency higher than a certain threshold, 5% by default, is accounted for, and the corresponding IUPAC ambiguity code is used. V-pipe also reports basic summary statistics, including the number of reads retained after quality control, the number of aligned reads, the region of the genome covered with a minimum number of aligned reads, and the minor allele frequencies per locus for all analysed samples.

We derive SNV calls from local haplotype reconstruction using ShoRAH (Zagordi *et al*., 2011). By considering co-occurring variants at multiple loci simultaneously, ShoRAH can effectively lower the SNV detection limit, i.e., the minimum frequency at which variants can be called reliably (McElroy *et al*., 2013). Alternatively, V-pipe also includes the variant caller LoFreq (version 2) (Wilm *et al*., 2012), which uses a position-wise and hence faster approach. A more complete characterization of the structure of a virus population consists in reconstructing the sequences and relative abundances of all viral haplotypes. For this purpose, we incorporate the quasispecies assembly tools HaploClique (Töpfer *et al*., 2014) and SAVAGE (Baaijens *et al*., 2017) to support reference-based and *de novo* haplotype reconstruction, respectively. Both of these methods are based on the overlap read graph, where reads correspond to nodes in the graph and edges indicate a sufficient overlap between corresponding reads. Conceptually, viral haplotypes are reconstructed iteratively by finding and merging max-cliques in the graph, i.e., locally reconstructed haplotypes are iteratively extended into full-length haplotypes.

As an additional feature, V-pipe includes a module to detect flow-cell cross contamination. To do so, reads are aligned to a panel of reference sequences containing suspected contaminants. We then detect and report the number of reads preferentially mapped to other references. The code developed to support this functionality, as well as various other steps of our pipeline, are maintained as an independent Python package called *smallgenomeutilities* (Supplementary Material Section S1.1).

V-pipe allows for constructing, maintaining, and using reproducible and traceable data analysis pipelines. In addition to providing a reproducible workflow and pinning dependency versions, V-pipe supports numerical reproducibility by fixing the seeds of all processing steps involving random sampling. Traceability is achieved by providing a well-documented pipeline in a well-defined framework and by ensuring meaningful log output in all processing steps. We have also made additional efforts to improve the reliability of individual tools used as part of our pipeline, such as ShoRAH and HaploClique (Supplementary Material Section S.1.2).

V-pipe is an open source project, the source code is freely available at https://github.com/cbg-ethz/V-pipe. Documentation including user guides can be found on the Wiki (https://github.com/cbg-ethz/V-pipe/wiki) and tutorials for specific viruses are available on our project page (https://cbg-ethz.github.io/V-pipe/). We also promote community involvement by maintaining a mailing list, providing user support, and offering workshops.

### 2.2 ngshmmalign: read aligner

Ngshmmalign performs a three-step alignment (Fig. 2). In the first step, reads are mapped to the reference genome to roughly determine their position. This initial mapping is done by indexing the reference genome using a *k*-mer index. To locate a read on the reference genome the *k*-mer index is queried by shifting a window of size *k* over the read. We then determine the mean and standard deviation of the returned location on the genome. If the standard deviation is above a certain threshold, the *k*-mer index match is considered suboptimal, and we perform a full genome-wide exhaustive alignment (Supplementary Material Section S1.3.2). After this initial mapping, reads are aligned in a semi-global mode using the Smith-Waterman algorithm. In the second step, the genome is partitioned into overlapping windows, and reads are assigned to windows based on the read-window overlaps. A multiple sequence alignment of the reads is performed independently for each of the windows by employing an iterative refinement approach implemented by the MAFFT software, specifically the L-INS-i method (Katoh and Standley, 2013) (Supplementary Material Section S1.3.3). We then infer the parameters of the profile HMM in a supervised manner, by assuming that the multiple read alignment represents a local sample of the profile HMM. In the third step, the final read alignment is obtained by re-aligning all reads to the profile HMM (Supplementary Material Section S1.3.4).

**Figure 2.**
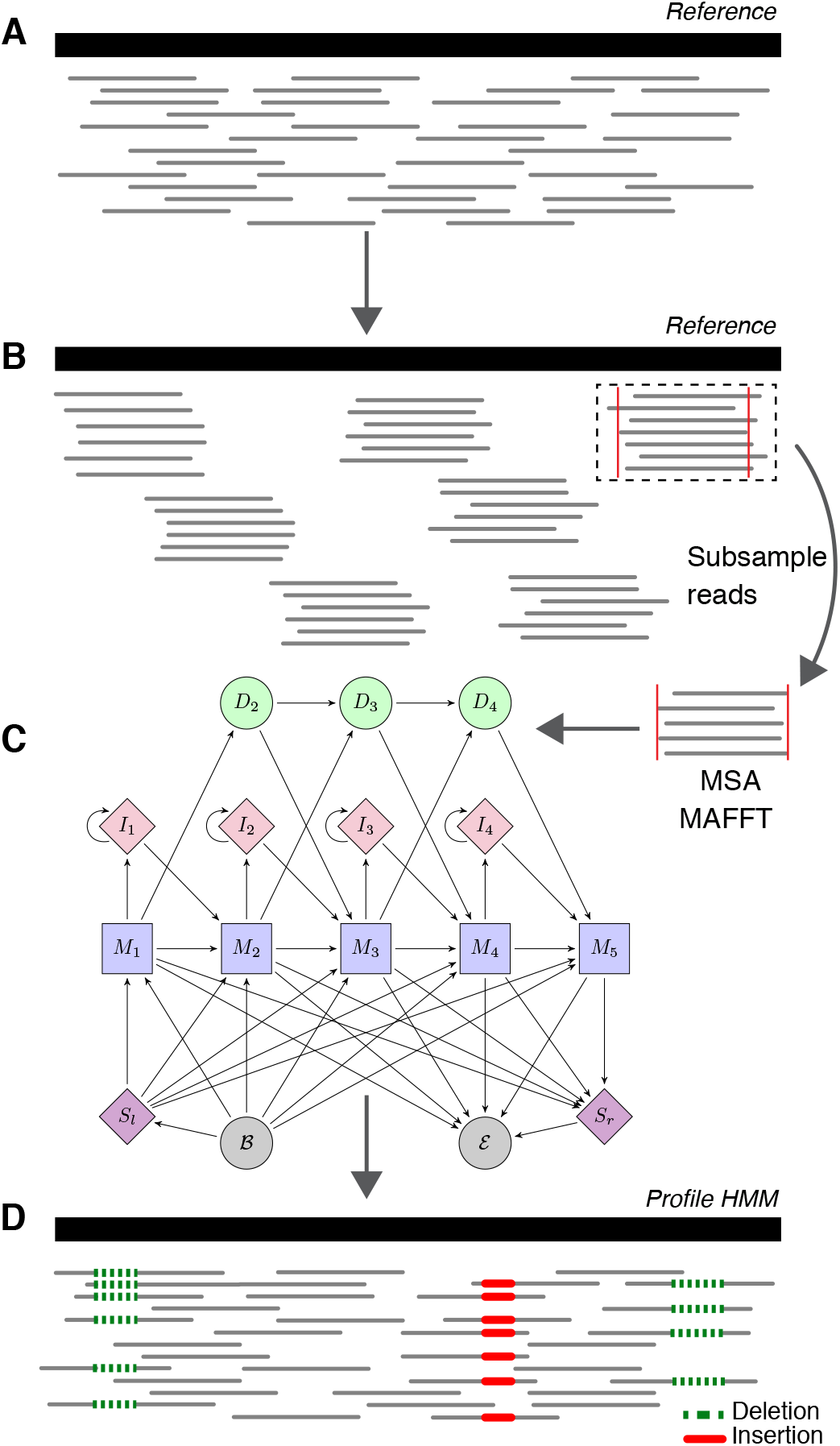
Multi-step strategy for read alignment. **A** Ngshmmalign uses a *k*-mer index for the initial approximate mapping of reads. If the *k*-mer index match is considered suboptimal, the alignment is done exhaustively. **B** Overlapping windows are extracted from the read mapping and a multiple sequence alignment is performed locally in each of the windows using the MAFFT software. **C** From the local multiple sequence alignments, the parameters of a global profile Hidden Markov Model (HMM) are estimated. **D** Reads are re-aligned against the profile HMM.

For every data set analysed within a single execution of the pipeline, ngshmmalign reports the read alignment using a position numbering relative to the profile. To standardise the position numbering, V-pipe performs a reference lift-over, i.e., it reports the alignment relative to a given reference sequence. To do so, we construct and use a multiple sequence alignment, containing the reference sequence and all consensus sequences from data sets included in the data analysis to transform the position numbering from the profile to the reference sequence. This multiple sequence alignment is also an intermediate result of V-pipe, which can be used for other applications, such as phylogenetic analyses.

### 2.3 Benchmarking

V-pipe provides benchmarking functionality by including two additional modules: (*i*) *simBench* can simulate sequencing reads from a mock virus population, and (*ii*) *testBench* can evaluate the accuracy of the inference results.

SimBench supports three modes. The first mode can be regarded as a simple quasispecies model (Domingo *et al*., 2005). A master sequence may be provided or it is generated by sampling *L* nucleotides uniformly at random, where L is the desired length of the sequence. Simulated haplotype sequences are generated from the master sequence by introducing substitutions, insertions, and deletions with user-configurable rates. Additionally, hyper-variable regions are simulated in the form of long deletions of variable length. In the second mode, the simulated haplotype sequences are sampled from a perfect binary tree. Every child node is generated from the parent sequence by introducing substitutions, insertions, and deletions with user-configurable rates. Here, we emulate explicitly the hierarchical evolutionary relationships among viral haplotypes. In the third mode, haplotype sequences are defined *a priori* and given as input to the pipeline, e.g., based on other models of viral evolution or on known viral sequences. The haplotype relative abundances are either (*i*) set to equal proportions (i.e. 1/n where *n* is the number of haplotypes), (*ii*) drawn from a Dirichlet distribution Dir(*α*) with concentration parameters *α_i_* = 1 for all haplotypes by default, or (*iii*) obtained using a geometric series with a given common ratio (0.75 by default).

After haplotype sequences have been generated, we use the ART software (Huang *et al*., 2012) to simulate either single-end or paired-end reads with configurable read length. ART is a read simulator with various built-in, technology-specific read error models and supports common sequencing errors, such as base substitutions, insertions, and deletions. We simulate reads from every individual haplotype with read coverage proportional to its relative abundance.

Instead of simulating sequencing reads, a user can also provide FASTQ files as input for the benchmark. This option allows for providing sequences simulated by different means, and it supports the analysis of control sequencing experiment with known haplotypes and relative abundances.

The evaluation module testBench is used to assess the accuracy of the inference results by comparing them to the ground truth. For SNV calling, we obtain a list of true SNVs by constructing a multiple sequence alignment of the underlying haplotypes and the reference sequence. Positions of expected SNVs are reported relative to the reference sequence. We then report the number of true positive, false positive, false negative, and true negative SNVs, as well as the inferred versus the expected SNV frequencies, the frequencies of false positives, and the number of false negatives per underlying haplotype.

V-pipe supports two modes for carrying out the benchmark, either evaluating a single pipeline arrangement at a time (*vpipeBench*), or multiple pipeline arrangements simultaneously (*vpipeBenchRunner*). Although the generation of synthetic data sets is reproducible, the latter mode should be preferred to ensure that other configuration settings are kept constant across different pipeline arrangements (i.e., combinations of processing steps).

### 2.4 Simulated data sets

We employ simBench to generate simulated reads, varying the number of haplotypes (ranging from 8 to 60), their relative abundances, and the total read coverage (either 10,000× or 40,000×, Supplementary Material Section S1.4). We use equal proportions for the haplotype abundances or sample their frequencies from a Dirichlet distribution with two sets of concentration parameters. We either use a symmetric Dirichlet distribution with *α_i_* = 1 for all *i* (denoted as Uniform), or choose one haplotype at random and assign to it a greater weight (*α*_0_ = 20 and *α_i_* = 1 for all *i* ≠ 0) to emulate a viral strain dominating in the population (denoted as Dirichlet). For all data sets, we simulate paired-end reads with 250 bp read length by using ART with the built-in quality profile for the MiSeq platform.

The data sets are based on HIV-1 or HCV sequences. Emulating populations using sequences derived from plasma samples of individual patients allows us to mimic the structure of viral populations more faithfully. Moreover, testing different viruses is crucial, as they may display different mechanisms of evolution and, hence, distinct forms of genetic variation. For the HIV-1-based data sets, we employ sequences of the HIV-1 subtype B envelope glycoprotein (*env*) gene obtained by single-genome amplification from individual patients (Lee *et al*., 2009). Particularly, we include in our simulated data sets sequences derived from subjects 1051 (GenBank accession numbers EU575134-EU575183) and BORI0637 (GenBank accession numbers EU576274-EU576302), for which 50 and 29 sequences are available, respectively. For the HCV data sets, we use 60 sequences (GenBank accession numbers KY565136-KY565195) from naturally occurring HCV genotype 1 subtype a (1a) E1E2 genes (El-Diwany *et al*., 2017).

As reference sequences, we use the HXB2 (GenBank accession number K03455.1) and H77 (Genbank accession number NC_004102.1) strain sequences, for HIV-1 and HCV, respectively.

### 2.5 Control sample for pipeline validation on real data sets

The control sample consists of an *in vitro* mixture of five known HIV-1 strains mixed at equal proportions. Four sequencing experiments were carried out starting with approximately 10^4^ (denoted 10K) and 10^5^ (denoted 100K) HIV-1 RNA copies. The samples were sequenced by using the Illumina MiSeq platform in paired-end read mode (2 × 250 bp length, v2 kit). The protocol described by Di Giallonardo *et al*. (2014) was employed for the amplification and sequencing. Primers were designed to cover almost the full HIV genome in five overlapping segments. Two types of sequencing experiments were carried out: one including all five amplicons (denoted A) and another one using only amplicon B (denoted B). This 2 × 2 design gives rise to four data sets referred to as A-10K, A-100K, B-10K, and B-100K.

## 3 Results

We demonstrate the benchmarking capabilities of V-pipe and evaluate the accuracy of SNV detection using different read aligners and mutation callers. In addition, we analyse an *in vitro* mixture of well-defined viral strains to assess the performance of one particular pipeline configuration on actual sequencing data. We also employ V-pipe to process publicly available longitudinal data from 11 patients (Zanini *et al*., 2015) to demonstrate the applicability of the pipeline as well as run time performance.

### 3.1 Simulation studies

We simulate reads from *in silico* mixtures of sequences derived from two different viruses, namely HIV-1 and HCV. We align simulated reads using ngshmmalign and compare it against two widely used read aligners, namely BWA MEM and Bowtie 2. For this comparison, we employ *vpipeBenchRunner* for the simultaneous execution of all pipeline configurations.

Averaged over all simulated data sets, the evaluated tools align more than 89% of the read pairs concordantly (Table S1). Although BWA MEM reports the highest percent of aligned reads, ngshmmalign aligns a larger portion of sequenced bases resulting in a higher average coverage (Fig. S1). To investigate potential bias in the alignment of reads due to differences in sequence similarity to the reference sequence, we report the fraction of aligned reads and aligned bases per haplotype. For the HCV-based data sets, sequences exhibit a broad range of divergence from the reference strain (0.005-0.112). BWA MEM aligns most of the reads regardless of the divergence from the reference, but a large fraction of the bases are soft-clipped, whereas ngshmmalign aligns a higher fraction of the bases for all haplotypes (Fig. S2).

Since the main focus of V-pipe is to infer viral genetic diversity, we evaluate the accuracy of the read aligners based on the F_1_ score of detecting SNVs using ShoRAH. The F_1_ score is the harmonic mean of precision and recall. To make the scores comparable, we report the performance metric for the union of all genomic loci covered by at least one of the aligners for each of the evaluated conditions. We evaluate two read coverages (10,000× and 40,000×) and three strategies to generate the underlying haplotype abundances. In most cases, ngshmmalign outperforms BWA MEM and Bowtie 2 in terms of the F_1_ score (Fig. 3A), and the difference in the scores is statistically significant (corrected p-value < 0.05, Wilcoxon signed-rank test, Supplementary Material Section S1.5.1). On the other hand, we do not observe substantial differences in the performance of the individual aligners while varying the distribution of haplotype frequencies, nor the evaluated coverages (Fig. S3).

**Figure 3.**
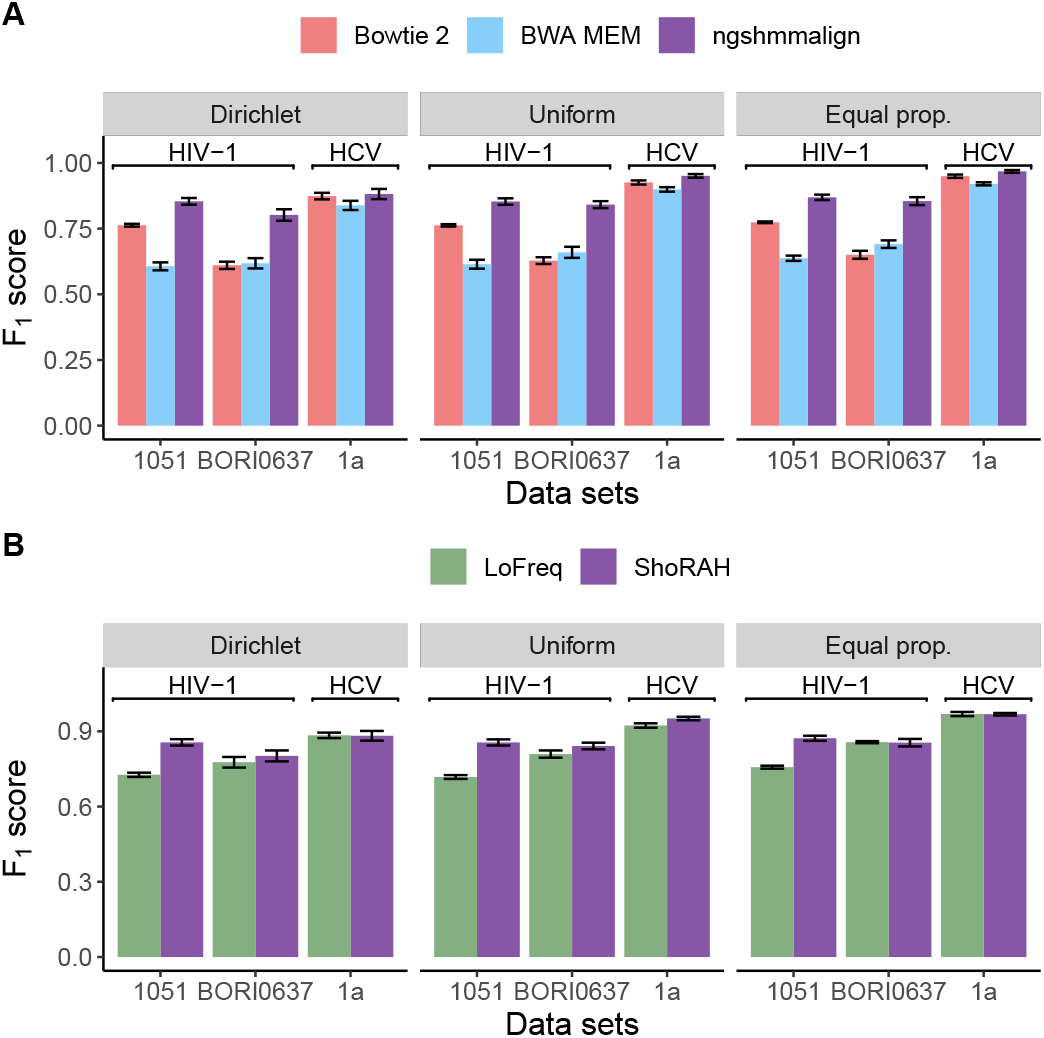
Performance of SNV detection on simulated data sets. **A** We compare ngshmmalign, BWA MEM and Bowtie 2 for read alignment, and fix ShoRAH for mutation calling. **B** We use ngshmmalign for the read alignment, and compare ShoRAH with LoFreq for mutation calling. In both panels, F_1_ scores are averaged over data sets with various numbers of haplotypes based on HIV-1 subtype B sequences from subjects 1051 and BORI0637, and HCV genotype 1a sequences. Results are shown for a read coverage of 10,000× and for different distributions of haplotype frequencies as described in the Methods Section 2.4 (Dirichlet: Dirichlet distribution with a high concentration parameter for one of the haplotypes (*α*_0_ = 20 and *α_i_* = 1 for all *i* ≠ 0). Uniform: symmetric Dirichlet distribution (*α_i_* = 1 for all *i*). Equal prop.: haplotype frequencies set to 1/*n* where *n* is the number of haplotypes). The error bar corresponds to the standard error.

Next, we focus on mutation calling and compare the accuracy of SNVs obtained by using ShoRAH versus LoFreq, while fixing ngshmmalign for the read alignment. We observe significant differences in the F_1_ scores for most data sets based on HIV-1 sequences from subject 1051 (corrected p-value < 0.05, Wilcoxon signed-rank test, Table S2); otherwise both tools appear to perform equally well (Figs. 3B and S4). Small discrepancies in the F_1_ scores can be attributed to differences in recall, whereas both tools show almost perfect precision (Figs. S5 and S6).

Although aligning reads with ngshmmalign and performing mutation calling with ShoRAH resulted in better F_1_ scores, there is a trade-off between accuracy and computational resources. On average and using 9 cores, ngshmmalign required 640 MB RAM and took 10 m 38 s, BWA MEM required 320 MB RAM and took 9 s, and Bowtie 2 required 332 MB RAM and took 25 s. For the mutation calling, ShoRAH took on average 50 m 47 s using 4.3 GB RAM and 9 cores, whereas LoFreq took on average 5 m 47 s using 75 MB RAM and executed as a single-threaded program. All the data sets were processed in 18-core Intel Xeon Gold 6140 processors.

### 3.2 Validation of V-Pipe on a mixture of five HIV-1 strains

A control sample is used to validate the performance of our pipeline on actual sequencing data, while full knowledge of the sample composition is still available.

specificity (Table 1). In data sets for which only amplicon B is sequenced (B-10k and B-100k), V-pipe reports perfect recall, whereas for data sets sequenced with all sets of primers (A-10k and A-100k), V-pipe misses a small fraction of the expected SNVs. The missed variants are predominately located at the genome termini, which correspond to regions of lower coverage. We also observe a decrease in precision for data sets A-100k, B-10k, and B-100k. Most of the falsely reported SNVs correspond to single-nucleotide deletions at very low frequencies, whereas such errors are less prominent in sample A-10k. When inspecting the precision of the mutation calls as a function of the variant frequencies, we find that a precision higher than 98% can be attained for SNVs with frequencies at least 0.5% for all the analysed data sets (Fig. 4A). In addition to detecting most of the expected variants, we find the frequencies of unique SNVs per haplotype reported by ShoRAH to be in good agreement with the relative abundances originally reported by Di Giallonardo *et al*. (2014) (Fig. 4B).

**Table 1.** Evaluating mutation calling using V-pipe. Data sets A and B result from using five overlapping amplicons (A) or only the second amplicon covering mainly HIV-1 pol (B), respectively, and the suffix indicates the initial amount of RNA copies We employ V-pipe for the data analysis using ngshmmalign with a *de novo* constructed reference sequence, and ShoRAH for SNV calling. In all cases, V-pipe detected more than 86% of the expected SNVs, with almost perfect

| Data set | Recall | Precision | Specificity |
| --- | --- | --- | --- |
| A-10k | 0.860 | 0.944 | 0.992 |
| A-100k | 0.873 | 0.631 | 0.925 |
| B-10k | 1 | 0.770 | 0.977 |
| B-100k | 1 | 0.403 | 0.885 |

**Figure 4.**
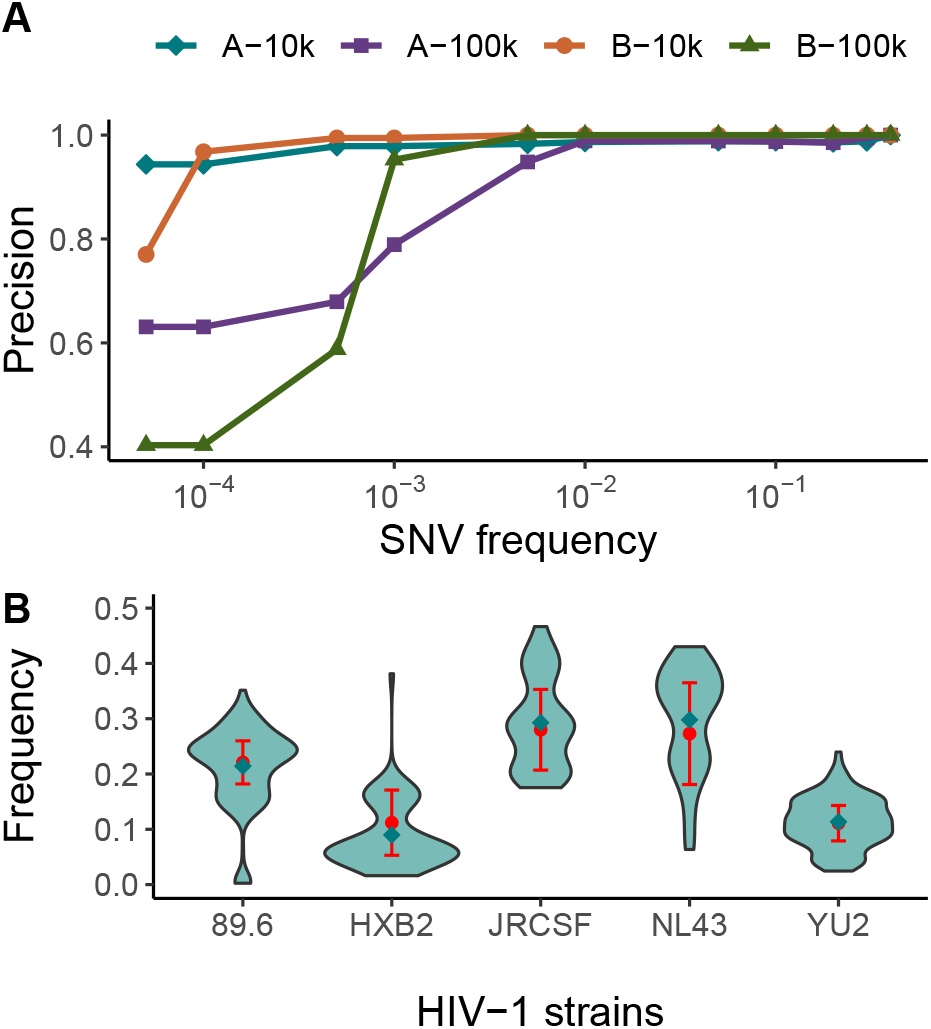
Evaluation of the performance of V-pipe on HTS data derived from the five-virus-mix. **A** Precision of SNV calls as a function of SNV frequency. **B** Distribution of inferred frequencies of unique SNVs for each of the five haplotype in the mix based on data set A-10k. The green diamonds show the corresponding average frequency. In red, we show the mean and standard deviation of estimated frequencies of reconstructed haplotypes from Illumina reads as originally reported by Di Giallonardo *et al*. (2014).

### 3.3 Application to clinical samples

To test V-pipe on clinical samples, we analyse publicly available data sets corresponding to longitudinal samples from one of the patients studied by Zanini *et al*. (2015) (ENA study accession number PRJEB9618). We employ V-pipe to infer local haplotypes on the 6 longitudinal data sets available for patient p2, using ngshmmalign and ShoRAH for local haplotype reconstruction. We compare the inferred haplotypes to the haplotype sequences reported by Zanini *et al*. (2015) on a region of the *p17* gene spanning nucleotides 872 to 1072 with respect to the HXB2 reference genome. This region has been arbitrarily chosen from regions displaying genetic diversity. After filtering out inferred haplotypes with a posterior probability smaller than 0.9 and an average read count of less than 10 reads, we find a perfect match between haplotype sequences reconstructed using V-pipe and the sequences previously reported by Zanini *et al*. (2015) using manual curated read alignments. However, V-pipe finds 1, 11, 3, and 8 additional haplotypes in the samples taken 561, 1255, 1628, and 2018 days after the estimated date of infection, respectively. The estimated frequencies of these haplotypes range from 0.08% to 6%. For the matching haplotypes, the absolute error in the estimated abundances is below 4% for all the considered data sets (Fig. S7). Monitoring viral genetic diversity over time provides an ideal basis for studying viral evolution. We visualise the reconstructed local haplotypes by representing pairwise Hamming distances in a two-dimensional plane (Fig. 5). We observe a drift in sequence space from the initial haplotype (in red, Fig. 5) as time progresses. Sequences of haplotypes recovered after 561 and 1255 days of infection are situated closer to the initial haplotype sequence, whereas sequences corresponding to later time points (after 1628 and 2018 days of infection) are further away.

**Figure 5.**
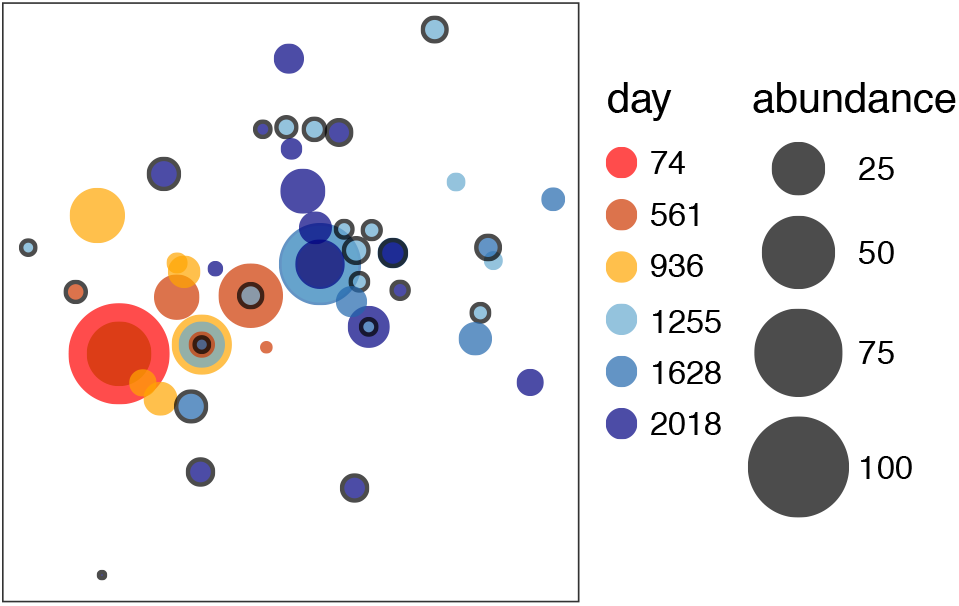
Representation of the reconstructed viral haplotypes in a region of the HIV-1 *p17* gene from longitudinal samples of patient p2 of Zanini *et al*. (2015). Discs represent inferred haplotypes, and their size reflect the relative abundances. Haplotypes are placed in a two-dimensional plane with the aim of preserving all pairwise Hamming distances as much as possible, using multi-dimensional scaling as implemented in scikit-learn (Pedregosa *et al*., 2011). The colors indicate the number of days after infection, and a circle around the disc denotes haplotypes identified by V-pipe but not by Zanini *et al*. (2015).

To illustrate the large-scale applicability of the pipeline, we note that using V-pipe to process all 92 data sets reported in Zanini *et al*. (2015) yielded an average run time per sample of 16 h on 12-core Intel Xeon E5-2680v3 processors (2.5-3.3 GHz), for an average coverage across samples of 31871×. Depending on the computational resources available, the total throughput can be largely independent of the number of samples, because the processing steps for individual data sets can be executed in parallel. Moreover, when computational resources are limited, V-pipe can be executed using alternative aligners, such as BWA MEM. For these particular data sets, BWA MEM aligned reads within 86 s on average, whereas ngshmmalign took on average 4 h 38 m. Similarly, LoFreq can be used for mutation calling as opposed to ShoRAH. In this case, ShoRAH took on average 11 h 7 m using 12 threads, whereas LoFreq took 2 h 42 m executed as a single-threaded program.

## 4 Discussion

Incorporating HTS technologies into viral genomics studies provides new opportunities to characterize intra-host virus populations in unprecedented depth. However, the applicability of these technologies is challenged by the amount and quality of the resulting data. In a clinical diagnostics setting, these large volumes of data need to be analysed accurately and efficiently within a short period of time.

To ensure reproducibility and traceability of the analysis workflows, we developed and implemented the bioinformatics pipeline V-pipe. V-pipe is tailored to studying intra-host diversity from viral HTS data. Although workflows for either the identification of minor variants (Huber *et al*., 2017) or the reconstruction of viral haplotypes (Jaya-sundara *et al*., 2015) have been proposed, to our knowledge, V-pipe is the first pipeline assessing viral diversity at all spatial levels of the viral genome, i.e., SNVs, as well as local and global haplotypes.

We also present a novel read alignment approach tailored to relatively small, yet highly diverse genomes. The aligner prioritises accuracy over speed and is based on profile hidden Markov models, which allow for capturing features shared among related sequences, such as mutations and indels, through position-specific probabilities.

We evaluated ngshmmalign on simulated reads from mock viral populations that attempt to mimic hyper-variable regions of HIV-1 (env gene) and HCV (*E1E2* region). The performance of the aligner was indirectly evaluated based on the detection of SNVs from the aligned reads. In most cases, the *F*_1_ score was found to be larger than 0.8, and SNV calls based on alignments obtained using ngshmmalign also reported a higher score compared to alignments by BWA MEM or Bowtie 2. Although ngshmmalign performed better under the evaluation conditions, it is possible that the alignments produced by BWA MEM or Bowtie 2 could be further improved by fine-tuning their hyper-parameters. Nevertheless, our results suggest that the SNV detection accuracy is to a larger extent influenced by the upstream alignment step than by the choice of the SNV caller. More importantly, with this simulation study, we demonstrated the benchmarking component implemented in V-pipe, which can be useful for adjusting the pipeline to meet specific requirements, e.g., on accuracy of the results or availability of computational resources.

Using V-pipe, we were able to reconstruct the genetic diversity present in a five-virus-mix control sample data set with high recall. While we observed lower precision, this could be improved substantially by filtering out deletion calls with frequencies below 0.5%. Choosing such thresholds for mutation and structural variant calling is highly dependent on the HTS protocols, but detecting SNVs at frequencies as low as 1% is already very close to the Illumina sequencing error rate.

In addition to simulated data sets and control sequencing samples, we demonstrated the applicability of the pipeline by processing data from 92 clinical samples resulting in typical run times of 16 h per sample. Thus, we believe that total turnaround times of roughly a few days are feasible using V-pipe’s default configuration (i.e., with ngshmmalign and ShoRAH) and including sample collection and preparation, sequencing, and data analysis. Hence, the accuracy as well as the run times of V-pipe make it suitable for many applications.

Most studies on viral genetic diversity have been limited to the identification of SNVs. Yet, the occurrence of mutations on the same genome might not be independent and the combined effect of co-occurring mutations might not be additive. On the other hand, it is not entirely obvious how to incorporate haplotype information, e.g., into monitoring epidemics, drug resistance surveillance, or supporting treatment decisions. There is, however, supporting evidence for using viral haplotypes to infer transmission pairs (Poon *et al*., 2016). V-pipe can support such research towards the understanding of the missing links in viral haplotyping.

In general, viral haplotype reconstruction is an active research field. Consequently, new computational methods for this task are regularly being proposed. In addition to novel methods, technological improvements are deployed at a high rate. Therefore, any pipeline needs to be actively maintained and constantly adapted to new requirements. We addressed this aspect from two directions. First, additional efforts have been directed into making the individual components of our pipeline, such as ShoRAH and HaploClique, more performant to handle current sequencing throughputs. Second, V-pipe features a modular and extensible architecture, such that the pipeline can be adapted to incorporate new tools. Furthermore, there is a pressing need to establish standards for data analysis. We thus introduced a benchmark component to support testing of different pipeline configurations and to contribute towards establishing best practices for viral bioinformatics.

One limitation of the pipeline is that it currently focuses on Illumina data. Other sequencing technologies such as Pacific Biosciences and Oxford Nanopore can produce longer reads which have the potential to reduce the complexity of the haplotype reconstruction problem. However the higher error rates, compared to Illumina sequencing, can be a limiting factor. Combining data from both short-read and long-read sequencing is a promising direction (Viehweger *et al*., 2019), and future developments should include extending the pipeline to support Pacific Biosciences and Oxford Nanopore data.

## Acknowledgments

We thank Tobias Marschall for outlining the needs of the community, contributing to the initial design of V-pipe, and co-organizing the 2017 Basel Computational Biology Conference (BC2) tutorial on “Production Pipelines for Virus Sequencing Data”. We thank Maryam Zaheri for testing, refactoring, and adding a continuous integration pipeline to the HaploClique software. We also thank Marek Pikulski and Nico Borgsmüller for critical reading.

