## Supplementary Material for "V-pipe: a computational pipeline for assessing viral genetic diversity from high-throughput sequencing data"

### 1 Supplementary material

#### S1.1 V-pipe: additional utilities

*smallgenomeutilities* is a collection of auxiliary scripts to support various analysis steps included in V-pipe. They allow, e.g., transforming the read alignment from one reference frame to another, constructing consensus sequences from aligned reads, obtaining frequencies of minor variants, providing statistics on the read coverage, and extracting regions of the read alignment with coverage above a certain threshold. The source code is available at https://github.com/cbg-ethz/smallgenomeutilities, and the package is also part of the PyPI and Bioconda (Grüning *et al.*, 2018) registries.

#### S1.2 Improvements of ShoRAH and HaploClique

We have made changes improving the reliability of two software tools integrated into V-pipe, namely ShoRAH and HaploClique. For ShoRAH, we ported the code to Python 3, modernized the build system, replaced expensive function calls with more efficient library functions and Intel SIMD intrinsics to boost computational efficiency, and implemented VCF file output. For HaploClique, we improved the build system, incorporated a continuous integration solution, and introduced functional modifications to reduce memory usage and execution times.

#### S1.3 Detailed description of ngshmmalign

Ngshmmalign is a novel sequence aligner developed to align HTS reads for small genomes. Many viruses are subject to high heterogeneity, stemming not only from single-nucleotide variants but also from structural variation. For instance, the HIV-1 *env* gene is the target of the immune system, and is therefore subject to strong positive selection. As a result, the locus is rather heterogeneous, with many newly inserted and deleted amino acids. A significant issue with current read aligners such as BWA (Li, 2013) and Bowtie (Langmead, 2012) is their reliance on index structures, which implicitly requires reads to have a very low edit distance to the reference genome. Furthermore, even if the seed region matches the reference genome, flanking indels are often aligned sub-optimally, as these aligners do not account for local structural heterogeneity.

Ngshmmalign performs a three-step alignment. In the first step, reads are mapped to the most likely location on the reference genome. In the second step, multiple read alignments are constructed locally, and a profile HMM is learned from them. The read alignment is then obtained in the third step by re-aligning all reads to the profile HMM, which can account for genomic structural variants found in the data. Below, each of the steps is described in more detail.

##### S1.3.1 Reading and checking FASTQ files

Ngshmmalign reads both single-end (e.g., 454, IonTorrent, Illumina) and paired-end (e.g., Illumina) FASTQ files. In the first step, the input is read and checked for consistency. This includes checking that no invalid bases are in the DNA string, the Phred score only contains values in the range from 0 to 40, and the length of the DNA string matches the Phred score string.

##### S1.3.2 Step 1: Read mapping

For performance reasons, the initial mapping is done with a $k$-mer index. The reference genome is indexed starting with $k = 20$. Ambiguous bases are expanded (e.g., R into A and G) into sequences of length $k$ comprised of only the bases A, C, G, and T. Unfortunately, in the worst case (i.e., a genome consisting of all N bases), this expansion can lead to all combinations in the full sequence space of $4^k$ sequences. If, in generating the $k$-mer index, the number of elements in the index exceeds a defined threshold of $10^6$ elements, then the index generation for $k$ is aborted and we proceed to build the $(k-1)$-mer index. This process is repeated until the total number of expanded sequences across the genome is below the threshold. In order to locate a read on the reference genome, the $k$-mer index is queried by shifting a window of size $k$ over the read.

The next step is to determine the mean and standard deviation of the returned location on the genome across the windows on the read. If the standard deviation is 0, a perfect linear match to a region on the reference has been found. If the standard deviation is non-zero but very small, then there is still very high confidence in the determined region. If the standard deviation is above a certain threshold (1000 by default), the location returned from the index is not trusted, and we perform a full genome-wide exhaustive alignment. The rationale of this approach is that we aim at detecting events that would result in reads mapping equally well to different locations of the reference genome, e.g., due to structural variants.

After mapping reads to a likely location on the genome, reads are aligned in a semi-global mode using the Smith-Waterman algorithm. The alignment of individual reads is performed in parallel.

##### S1.3.3 Step 2: Estimating profile HMM parameters

After the initial mapping, we know the approximate location and strandedness of the HTS reads on the genome. The genome is then partitioned into overlapping windows. The size of the window is $\frac{6}{5}$ of the average read length, and adjacent windows are shifted by $\frac{1}{5}$ of the average read length. For example, for 300 nt reads, the window size would be 360 nt on the reference genome, and windows are shifted by 60 nt. In effect, the largest overlap between non-identical windows is equal to the read length. The reads are binned into the windows based on the overlap between read and window. Each read fits completely into one window. The special case, in which a read falls exactly into the overlap of two windows is resolved by selecting one of the two windows uniformly at random. For every window, we sample without replacement 500 reads among those that have at least 80% of their bases aligned to the genome. This is done to limit the computational resources required for the subsequent multiple sequence alignment step. This step is performed independently for each of the windows by employing an iterative refinement approach implemented by the MAFFT software, specifically the L-INS-i method (Katoh and Standley, 2013).

After all multiple sequence alignments have been performed, the parameters of the profile HMM are inferred. This is done by assuming that the multiple sequence alignments locally represent the profile HMM, such that the parameters can be learned in a supervised manner. The match-to-match, match-to-deletion, deletion-to-match, and deletion-to-deletion transition probabilities of the hidden Markov model are estimated from the individual window-wise alignments. If the match-to-deletion frequency lies below a certain threshold (5% by default), then the absence of a biological deletion is assumed and the value is set to the technical sequencing match-to-deletion probability. The same argument is applied to deletion-to-deletion transitions, which are set to technical sequencing error probabilities if the deletion-to-match frequencies are below the 5% threshold. A position where at least 5% of reads have a base, as opposed to a gap in the multiple sequence alignment, is considered a biologically relevant position and is added to the profile. This leads to a situation where true biological insertions are included into the profile, such that ideally all remaining insertions are technical errors. This is desirable for downstream analyses, as modeling insertions is generally harder than modeling deletions. For instance, most reference-based haplotype reconstruction tools ignore insertions, but account for deletions.

##### S1.3.4 Step 3: Alignment of reads to the profile HMM

After inferring the profile HMM parameters, reads are aligned against the profile. For performance reasons, we assume that the strandedness of the reads inferred in the mapping step is correct. All reads are aligned in a fully concurrent fashion by processing 500 reads per thread by default.

A limitation with other read mappers, such as BWA and Bowtie, is the deterministic placement of small deletions in homopolymeric regions, usually by placing them at the beginning of the homopolymer. These deletions likely correspond to technical errors due to polymerase slippage and can result in falsely reporting as variants spurious artefacts in downstream variant calling steps. Ngshmmalign addresses this issue by determining all co-optimal alignments and creating a per-site deletion histogram. For every read, the alignment is compared to this deletion histogram, and the final alignment is chosen such that deletions are evenly distributed across the homopolymeric region. This avoids choosing co-optimal alignments with many deletions aligning at the same position by chance.

##### S1.3.5 Output of ngshmmalign

Ngshmmalign further checks for the proper orientation of paired-end reads. Paired-end reads, for which both mates align in forward or reverse direction, are considered invalid and written to a separate file. Finally, reads are sorted first by their identifier and then by position, before being written out in SAM format. Ngshmmalign outputs the alignment as a fully compliant SAM file that passes all filters of picard (, http://broadinstitute.github.io/picard). The profile HMM is also output in a human-readable format that can be edited by users to adjust parameters in order to perform additional alignment iterations. Besides the SAM output file, ngshmmalign produces both the majority-vote and the ambiguity-coded consensus sequences. We use lowercase characters in the consensus sequences to distinguish positions at which no or insignificant coverage was encountered, such that users can filter such positions.

#### S1.4 Additional details on simulated data sets

The number of haplotypes in each constructed population ranges from 8 to 60. For the HIV-1-based data sets, we subsample sequences from a given patient to generate data sets with 12, 25, and 50 haplotypes from subject 1051, and with 8, 15, 28, and 29 haplotypes from subject BORI0637. For the HCV-based data sets, we subsample sequences from different genotypes to generate data sets with 8, 15, 30, and 60 haplotypes from genotype 1a. Haplotypes were mixed either at equal proportions or by drawing their relative frequencies from a Dirichlet distribution. We evaluate two strategies for the concentration parameters of the Dirichlet distribution. We either use a symmetric distribution with $\alpha_i = 1$ for all $i$, or a modification thereof obtained by choosing one haplotype $j$ at random and setting its weight to $\alpha_j = 20$. The former is equivalent to a uniform distribution over the standard $(n-1)$-simplex. The latter is used to emulate a single viral strain dominating the population, while other strains co-exist at lower abundances. For each population, we generate data sets with read coverages of 10,000$\times$ and 40,000$\times$. For each combination of number of haplotypes, read coverage, and haplotype frequency distributions, we simulate 6 independent data sets.

#### S1.5 Detailed benckmarking results

We demonstrate V-pipe's benchmarking functionality by assessing the accuracy of SNV detection using various read aligners and mutation callers.

**Table S1.** Fraction of aligned reads averaged across all simulated data sets for each given virus type

| Virus | ngshmmalign | BWA MEM | Bowtie |
|---|---|---|---|
| HIV-1 | 0.966 | 0.998 | 0.955 |
| HCV | 0.985 | 0.999 | 0.892 |

The results summarised in Figure 3 panels A and B are shown in more detail in Figures S3 and S4 , respectively. The low $F_1$ scores observed for the HIV-1-based data sets with 29 haplotypes (below 0.5 in most cases, Figs. S3 and S4 ) are attributed to the fact that one viral sequence from subject BORI0637 exhibits two very large deletions (around 230 bp each), which are almost as long as the read length. This particular sequence was excluded from the remaining data sets. In addition, the data sets containing this sequence were not accounted for in the main text figures.

##### S1.5.1 Statistical test

We use the Wilcoxon signed-rank test to assess the difference in the $F_1$ scores while grouping the results by the read coverage and the strategy used to generate the haplotype abundances. In most cases, the differences in the performance metric between ngshmmalign and each of the competitors remain significant at a 5% significance level after correcting for multiple comparisons using the Bonferroni correction method (Table S2).

**Table S2.** Evaluation of the differences in $F_1$ scores of SNV calls using the Wilcoxon signed rank test. The reported p-values are corrected using the Bonferroni correction method. Dirichlet: haplotype frequencies sampled from a Dirichlet distribution with a high concentration parameter for one of the haplotypes ($\alpha_0 = 20$ and $\alpha_i = 1,\ i \neq 0$). Uniform: haplotype frequencies sampled from a symmetric Dirichlet distribution ($\alpha_i = 1, \forall i$). Equal prop.: haplotype frequencies are set to $1/n$ where $n$ is the number of haplotypes

| Data sets | Compared tools | Coverage | Haplotype distribution | p-value |
|---|---|---|---|---|
| HIV-1 / 1051 | ngshmmalign vs BWA MEM | all | all | $1.194 \times 10^{-17}$ |
| | | 10000× | Dirichlet | $4.807 \times 10^{-4}$ |
| | | 10000× | Uniform | $4.807 \times 10^{-4}$ |
| | | 10000× | Equal proportions | $4.807 \times 10^{-4}$ |
| | | 40000× | Dirichlet | $4.807 \times 10^{-4}$ |
| | | 40000× | Uniform | $4.807 \times 10^{-4}$ |
| | | 40000× | Equal proportions | $4.807 \times 10^{-4}$ |
| | ngshmmalign vs Bowtie 2 | all | all | $1.262 \times 10^{-17}$ |
| | | 10000× | Dirichlet | $4.807 \times 10^{-4}$ |
| | | 10000× | Uniform | $4.807 \times 10^{-4}$ |
| | | 10000× | Equal proportions | $4.807 \times 10^{-4}$ |
| | | 40000× | Dirichlet | $4.807 \times 10^{-4}$ |
| | | 40000× | Uniform | $9.613 \times 10^{-4}$ |
| | | 40000× | Equal proportions | $4.807 \times 10^{-4}$ |
| HIV-1 / BORI0637 | ngshmmalign vs BWA MEM | all | all | $1.846 \times 10^{-22}$ |
| | | 10000× | Dirichlet | $3.755 \times 10^{-5}$ |
| | | 10000× | Uniform | $7.510 \times 10^{-6}$ |
| | | 10000× | Equal proportions | $5.257 \times 10^{-5}$ |
| | | 40000× | Dirichlet | $8.261 \times 10^{-4}$ |
| | | 40000× | Uniform | $7.510 \times 10^{-5}$ |
| | | 40000× | Equal proportions | $7.510 \times 10^{-6}$ |
| | ngshmmalign vs Bowtie 2 | all | all | $2.491 \times 10^{-23}$ |
| | | 10000× | Dirichlet | $7.510 \times 10^{-5}$ |
| | | 10000× | Uniform | $1.502 \times 10^{-5}$ |
| | | 10000× | Equal proportions | $7.510 \times 10^{-6}$ |
| | | 40000× | Dirichlet | $5.257 \times 10^{-5}$ |
| | | 40000× | Uniform | $7.510 \times 10^{-6}$ |
| | | 40000× | Equal proportions | $7.510 \times 10^{-6}$ |

**Table S2.** Evaluation of the differences in $F_1$ scores of SNV calls using the Wilcoxon signed rank test. The reported p-values are corrected using the Bonferroni correction method. Dirichlet: haplotype frequencies sampled from a Dirichlet distribution with a high concentration parameter for one of the haplotypes ($\alpha_0 = 20$ and $\alpha_i = 1,\ i \neq 0$). Uniform: haplotype frequencies sampled from a symmetric Dirichlet distribution ($\alpha_i = 1,\ \forall i$). Equal prop.: haplotype frequencies are set to $1/n$ where $n$ is the number of haplotypes

| Data sets | Compared tools | Coverage | Haplotype distribution | p-value |
|---|---|---|---|---|
| HCV | ngshmmalign vs BWA MEM | all | all | $1.804 \times 10^{-20}$ |
| | | 10000× | Dirichlet | $3.357 \times 10^{-3}$ |
| | | 10000× | Uniform | $7.510 \times 10^{-6}$ |
| | | 10000× | Equal proportions | $7.510 \times 10^{-6}$ |
| | | 40000× | Dirichlet | 0.04677 |
| | | 40000× | Uniform | $7.510 \times 10^{-6}$ |
| | | 40000× | Equal proportions | $5.163 \times 10^{-6}$ |
| | ngshmmalign vs Bowtie 2 | all | all | $1.958 \times 10^{-13}$ |
| | | 10000× | Dirichlet | 1 |
| | | 10000× | Uniform | $7.510 \times 10^{-6}$ |
| | | 10000× | Equal proportions | $1.900 \times 10^{-3}$ |
| | | 40000× | Dirichlet | 1 |
| | | 40000× | Uniform | $1.427 \times 10^{-4}$ |
| | | 40000× | Equal proportions | $3.357 \times 10^{-3}$ |
| HIV-1 / 1051 | ShoRAH vs LoFreq | all | all | $2.463 \times 10^{-17}$ |
| | | 10000× | Dirichlet | $4.807 \times 10^{-4}$ |
| | | 10000× | Uniform | $4.807 \times 10^{-4}$ |
| | | 10000× | Equal proportions | $4.807 \times 10^{-4}$ |
| | | 40000× | Dirichlet | $4.807 \times 10^{-4}$ |
| | | 40000× | Uniform | 0.05287 |
| | | 40000× | Equal proportions | $4.807 \times 10^{-4}$ |
| HIV-1 / BORI0637 | ShoRAH vs LoFreq | all | all | $7.627 \times 10^{-3}$ |
| | | 10000× | Dirichlet | 0.198 |
| | | 10000× | Uniform | 0.499 |
| | | 10000× | Equal proportions | 1 |
| | | 40000× | Dirichlet | 1 |
| | | 40000× | Uniform | 1 |
| | | 40000× | Equal proportions | 1 |
| HCV | ShoRAH vs LoFreq | all | all | 0.0252 |
| | | 10000× | Dirichlet | 1 |
| | | 10000× | Uniform | $3.755 \times 10^{-5}$ |
| | | 10000× | Equal proportions | 1 |
| | | 40000× | Dirichlet | 1 |
| | | 40000× | Uniform | $8.006 \times 10^{-3}$ |
| | | 40000× | Equal proportions | 1 |

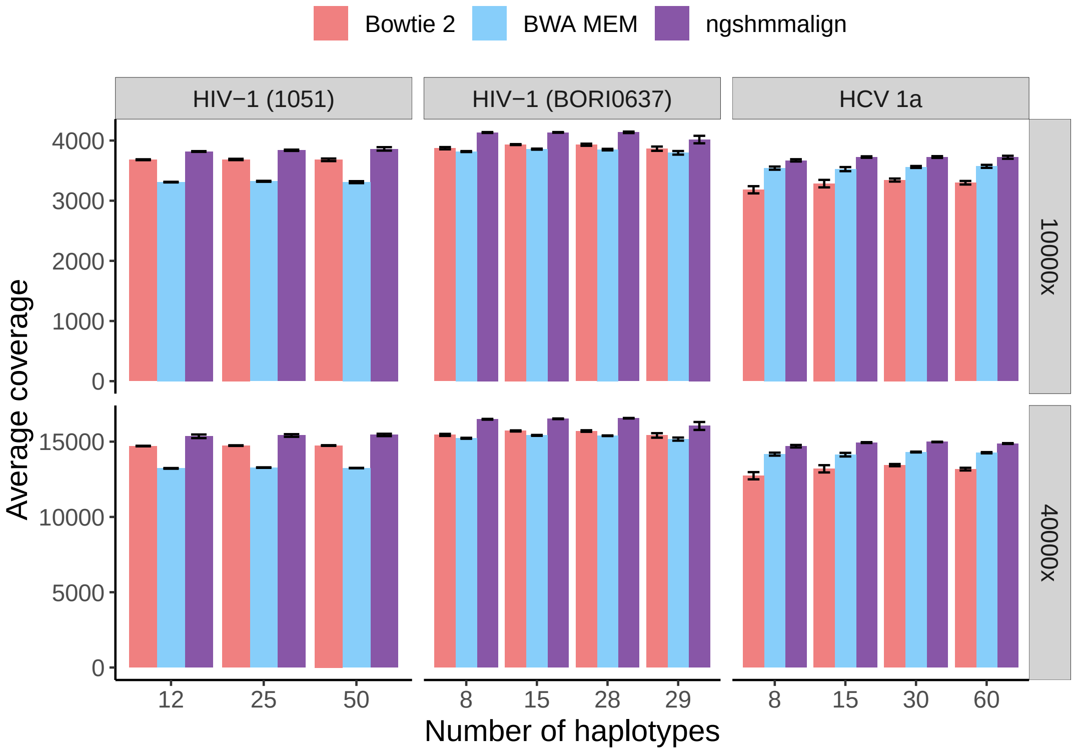


**Figure S1 .** Average read coverage after read alignment for simulated data sets with different numbers of haplotypes. The upper and lower panel show results for the corresponding initial read coverage (i.e., before QC and read alignment). The colored bars depict results obtained by using different read aligners, namely Bowtie 2, BWA MEM and ngshmmalign. The error bar corresponds to the standard error across 18 data sets with same number of underlying haplotypes and initial read coverage, but different distributions of the haplotype frequencies.

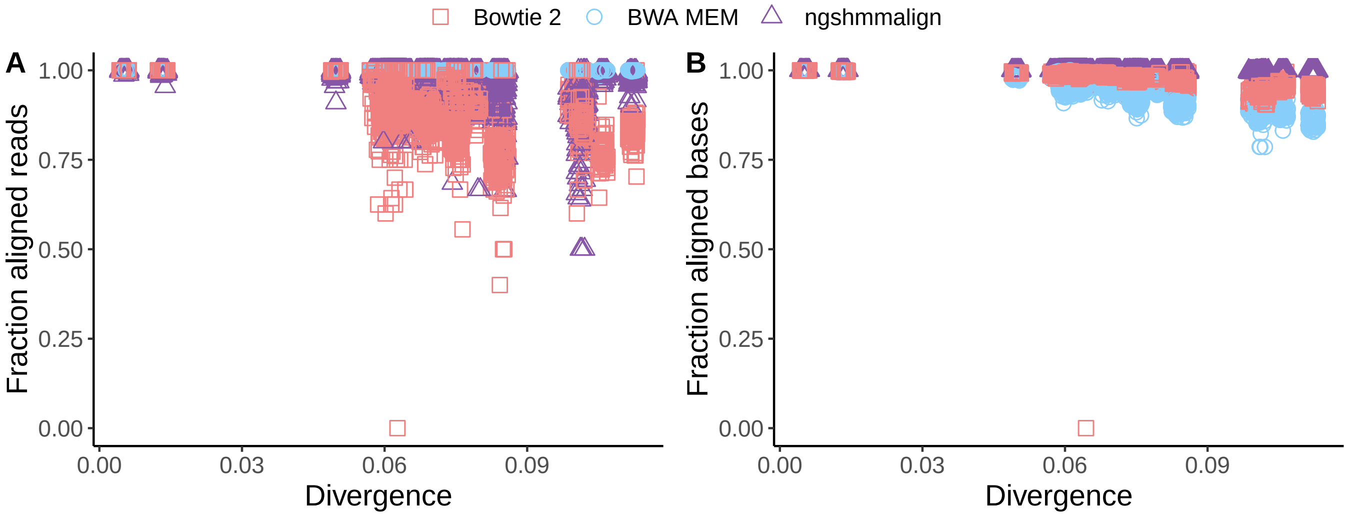


**Figure S2 .** Evaluation of potential alignment bias due to differences in divergence between the underlying haplotypes and the reference strain. We plot the fraction of **A** aligned reads and **B** aligned bases versus the divergence from the reference strain. Divergence is estimated as the Hamming distance between each haplotype and the reference strain divided by the sequence length (x-axis). The colored shapes depict results obtained by aligning reads with Bowtie 2, BWA MEM, or ngshmmalign.

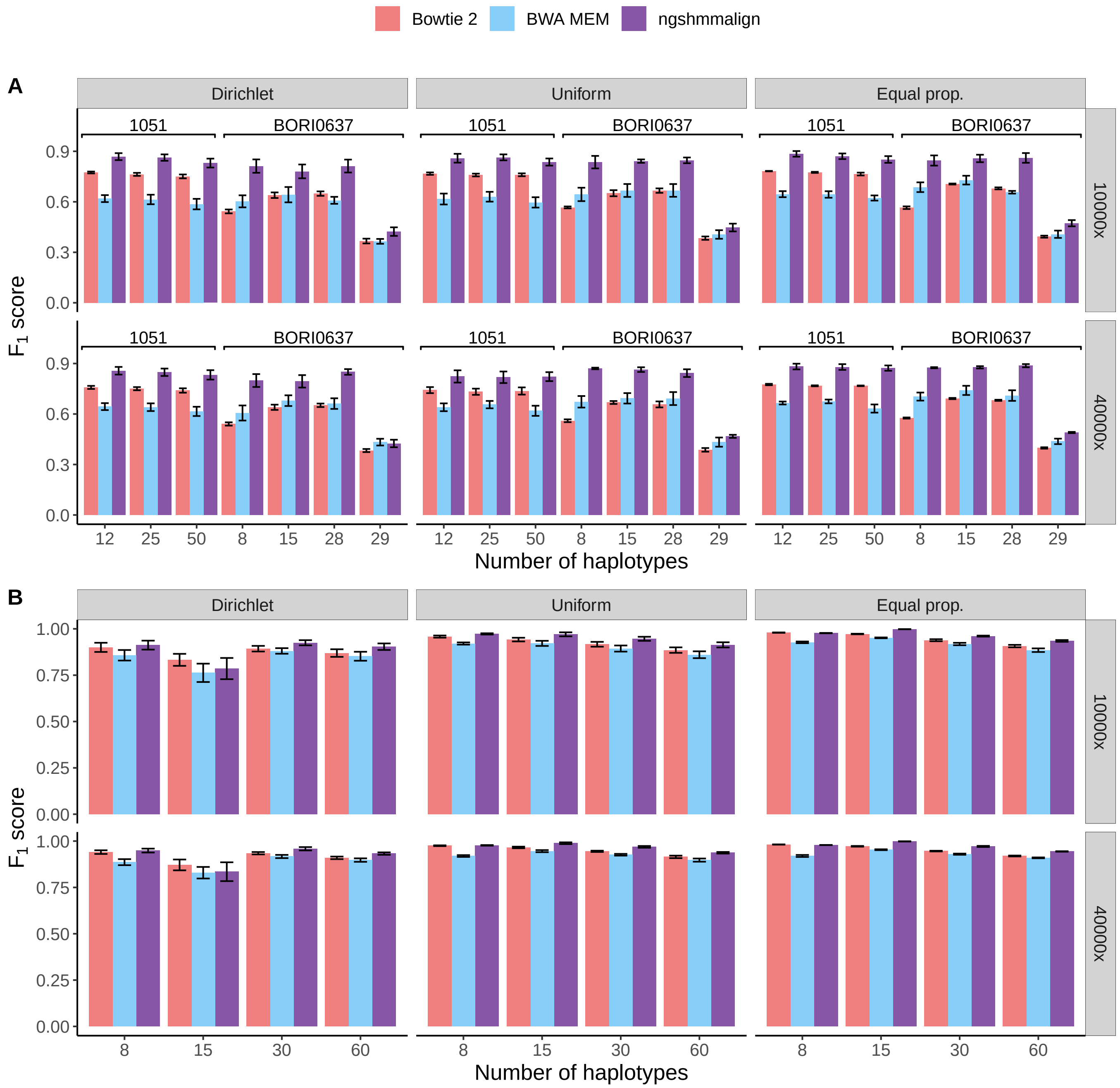


**Figure S3 .** Performance of SNV detection on simulated data sets for different read aligners. We use ngshmmalign, BWA MEM or Bowtie 2 for the read alignment, and fix ShoRAH for mutation calling. We simulate data sets with different number of haplotypes, read coverages and distributions of haplotype frequencies. $F_1$ scores of SNV calls are shown for data sets based on **A** HIV-1 sequences from subjects 1051 and BORI0637, and on **B** HCV genotype 1a sequences. The error bar corresponds to the standard error across 6 replicates. Dirichlet: haplotype frequencies sampled from a Dirichlet distribution with a high concentration parameter for one of the haplotypes ($\alpha_0 = 20$ and $\alpha_i = 1$, $i \neq 0$). Uniform: haplotype frequencies sampled from a symmetric Dirichlet distribution ($\alpha_i = 1$ for all $i$). Equal prop.: haplotype frequencies are set to $1/n$ where $n$ is the number of haplotypes

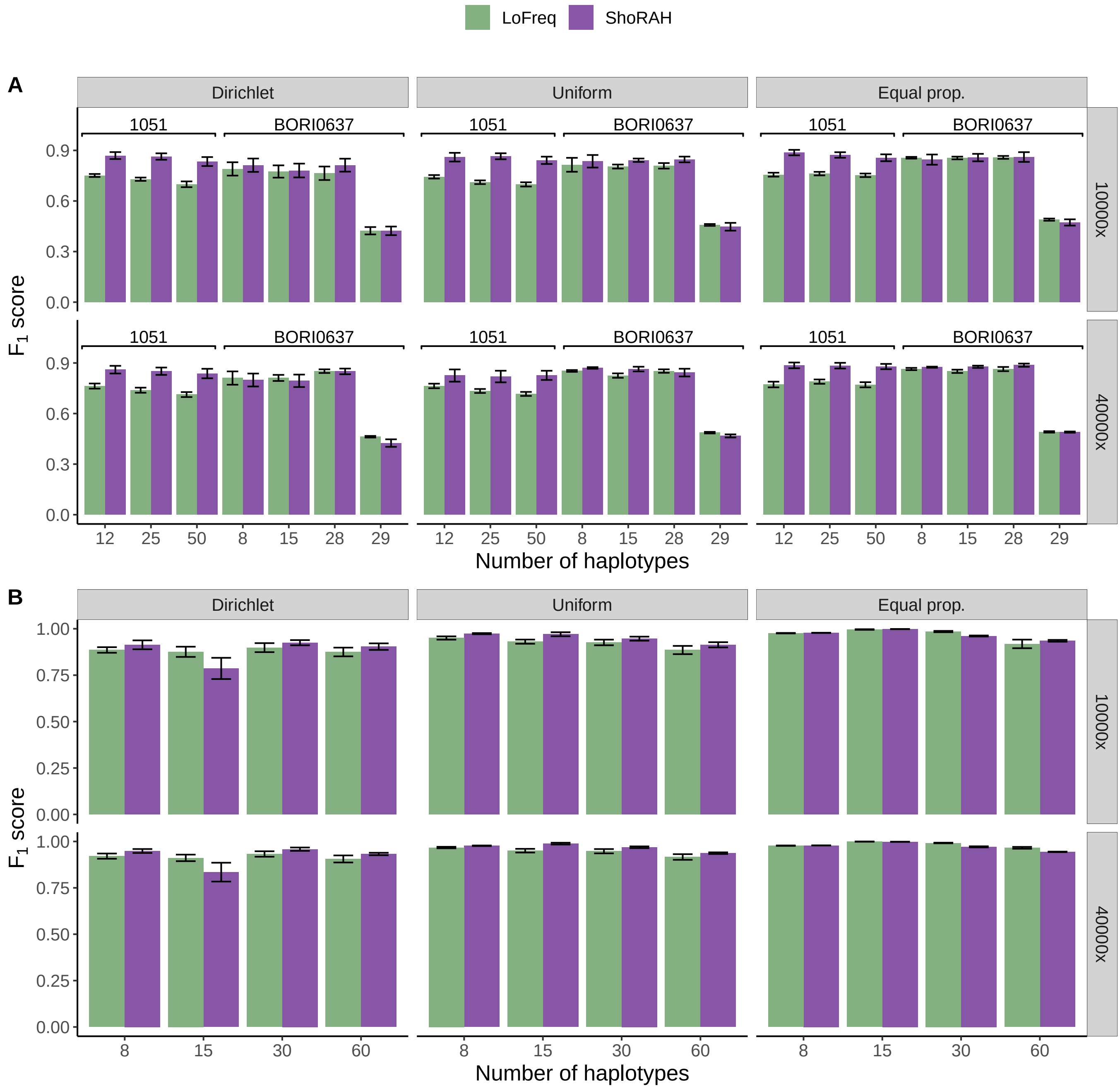


**Figure S4 .** Evaluation of the performance of different tools for SNV calling on simulated data sets. We fix ngshmmalign for the read alignment, and compare ShoRAH with LoFreq for mutation calling. We simulate data sets with different number of haplotypes, different read coverages and distributions of haplotype frequencies. **A** $F_1$ scores of SNV calls are shown for data sets based on HIV-1 sequences from subjects 1051 and BORI0637. **B** $F_1$ scores of SNV calls are shown for data sets based on HCV genotype 1a sequences. The error bar corresponds to the standard error across 6 replicates. Dirichlet: haplotype frequencies sampled from a Dirichlet distribution with a high concentration parameter for one of the haplotypes ($\alpha_0 = 20$ and $\alpha_i = 1$, $i \neq 0$). Uniform: haplotype frequencies sampled from a symmetric Dirichlet distribution ($\alpha_i = 1$ for all $i$). Equal prop.: haplotype frequencies are set to $1/n$ where $n$ is the number of haplotypes

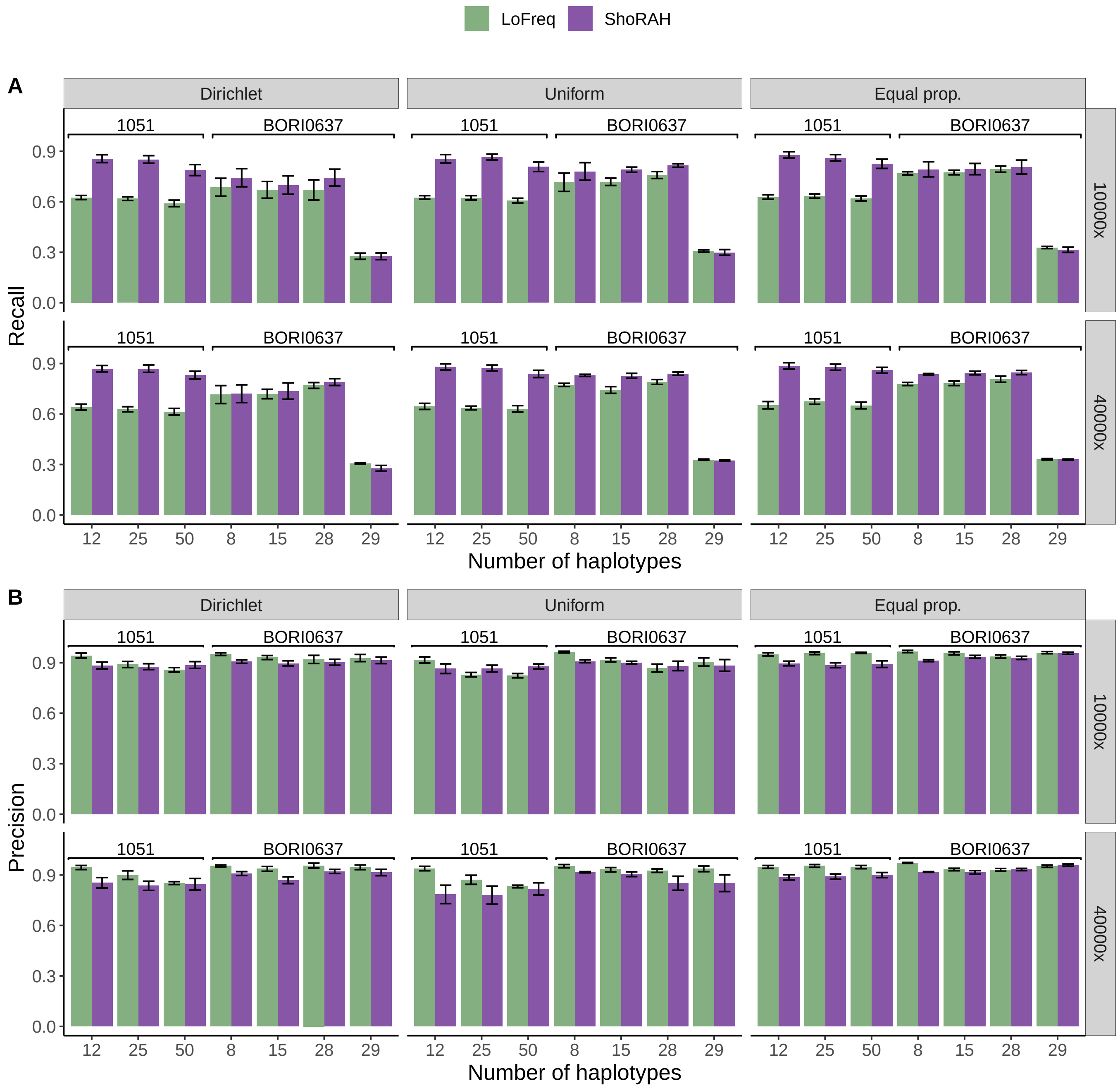


**Figure S5 .** Evaluation of the performance of different tools for SNV calling on HIV-1-based simulated data sets. We fix ngshmmalign for the read alignment, and compare ShoRAH with LoFreq for mutation calling. **A** Recall and **B** precision of SNV calls as a function of number of haplotypes, and for different read coverages and distributions of haplotype frequencies. The error bar corresponds to the standard error across 6 replicates. Dirichlet: haplotype frequencies sampled from a Dirichlet distribution with a high concentration parameter for one of the haplotypes ($\alpha_0 = 20$ and $\alpha_i = 1,\ i \neq 0$). Uniform: haplotype frequencies sampled from a symmetric Dirichlet distribution ($\alpha_i = 1$ for all $i$). Equal prop.: haplotype frequencies are set to $1/n$ where $n$ is the number of haplotypes

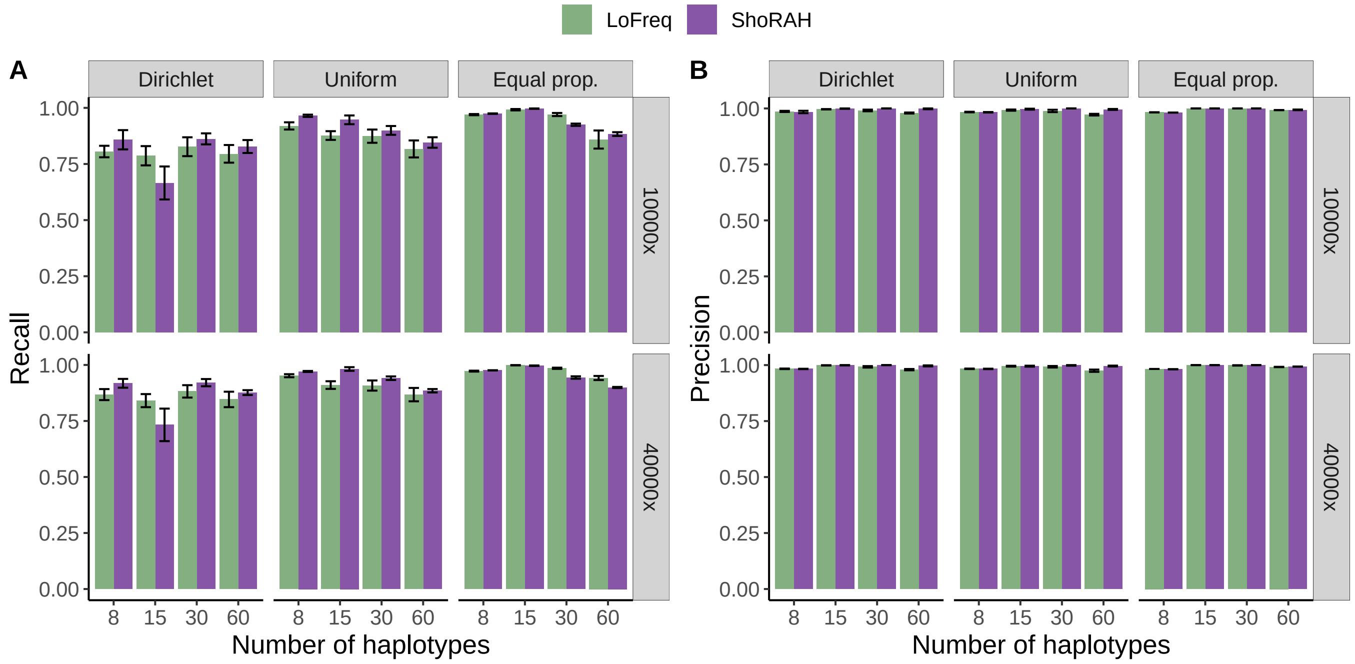


**Figure S6 .** Performance of SNV detection for different mutation callers on HCV-based simulated data sets. We fix ngshmmalign for the read alignment, and compare ShoRAH with LoFreq for mutation calling. **A** Recall and **B** precision of SNV calls as a function of number of haplotypes, and for different read coverages and distributions of haplotype frequencies. The error bar corresponds to the standard error across 6 replicates. Dirichlet: haplotype frequencies sampled from a Dirichlet distribution with a high concentration parameter for one of the haplotypes ($\alpha_0 = 20$ and $\alpha_i = 1,\ i \neq 0$). Uniform: haplotype frequencies sampled from a symmetric Dirichlet distribution ($\alpha_i = 1$ for all $i$). Equal prop.: haplotype frequencies are set to $1/n$ where $n$ is the number of haplotypes.

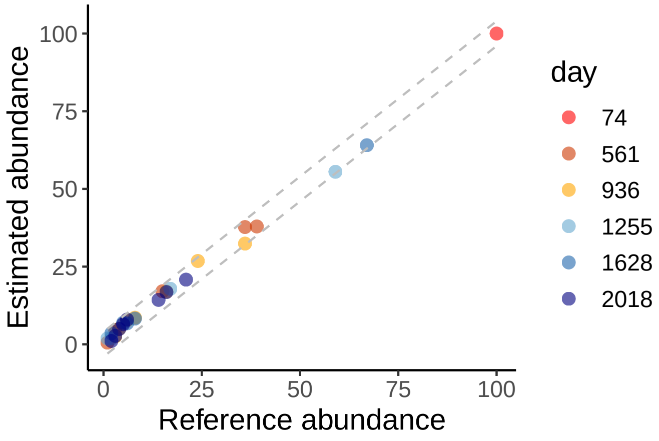


**Figure S7 .** Comparison of estimated versus reported haplotype abundances from longitudinal samples of patient p2 of Zanini *et al.* (2015). Dotted lines delimit a $\pm 4\%$ abundance error band centered around the previously reported abundances.
